# Short Neuropeptide F regulates the starvation mediated enhanced locomotor Activity in *Drosophila*

**DOI:** 10.1101/764688

**Authors:** Anna Geo, Himani Pathak, Anamika Elizabeth Kujur, Sreesha R Sudhakar, Nisha N Kannan

**Affiliations:** Chronobiology Laboratory, School of Biology, Indian Institute of Science Education and Research (IISER), Thiruvananthapuram, Kerala 695551, India; Drosophila Energy and Metabolism Laboratory, School of Biology, Indian Institute of Science Education and Research (IISER), Thiruvananthapuram, Kerala 695551, India

**Keywords:** circadian, short neuropeptide F, hyperactivity, oscillation, starvation, triglyceride metabolism

## Abstract

The circadian clock regulates various behavioral, metabolic and physiological processes to occur at the most suitable time of the day. Internal energy stores and nutrient availability modulates the most apparent circadian clock mediated locmotor activity rhythm in *Drosophila*. Although previous studies unraveled the role of circadian clock in metabolism and activity rest rhythm, the precise pathway through which the circadian neuropeptidergic signaling regulates internal energy storage and the starvation-mediated increase in activity resembling foraging remains largely unclear. This study was aimed to elucidate the role of circadian neuropeptide, short neuropeptide F (sNPF) in triglyceride metabolism, starvation resistance and starvation-mediated increased locomotor activity in *Drosophila*. The results showed that *snpf* transcripts exhibits significant rhythmicity in wild type flies under 12:12 hour light-dark cycles (LD) and constant darkness (DD) whereas *snpf* transcript level in *period* null flies did not exhibit any significant rhythmicity under LD. Knockdown of sNPF in circadian clock neurons reduced the triglyceride level, starvation resistance and increased the starvation-mediated hyperactivity response after 24 hour of starvation. Further studies showed that knock down of sNPF receptors (sNPFR) expressed in insulin producing cells (IPC) increased the starvation resistance and reduced starvation-induced hyperactivity response after 24 hour of starvation. Collectively, our results suggest that transcriptional oscillation of *snpf* mRNA is endogenously controlled by the circadian clock and elucidate the role of sNPF in modulating locomotor activity in accordance with the nutrient availability in *Drosophila*.

## Introduction

The circadian clock drives daily rhythms in a wide array of physiological and behavioral processes by scheduling it at the appropriate time of the day. This inherent timekeeping system coordinates the phase of two fundamental behavioral rhythms such as feeding and sleep/wake cycles and it is believed to aid the organism to adapt with the cyclic external environmental changes. The growing body of evidence indicates that the circadian clock not only mediates optimal phasing of behavioral rhythmicity during 24 hours of a day but also intimately ties with metabolism (Eckel-Mahan and Sassone-Corsi, 2013). While the clock orchestrates numerous metabolic pathways, the nutrient availability and metabolic status can in turn feed back to impinge on the functioning of the circadian clock (Damiola et al., 2000; Nakahata et al., 2009; Vollmers et al., 2009; Xu et al., 2011). A large body of evidence accumulated over the past two decades remarkably improved our understanding of the essential role of the clock in metabolism, feeding behavior, and the sleep wake cycle (Maury et al., 2010). However, little is known about the precise underlying pathways by which the circadian clock aids the organism to attain temporal harmonics between the sleep wake cycle and feeding behavior.

The association between the circadian clock, sleep wake cycle and metabolism was further substantiated by the findings that the circadian timing system drives rhythmicity of various peptides involved in metabolism that also play key role in the regulation of sleep wake cycles (Reviewed in Kalsbeek and Fliers, 2013). For instance, peptides involved in metabolism such as leptin, ghrelin and cholecystokinin display circadian rhythmicity in expression (Ahima et al., 1998; Kalra et al., 2003; Kalsbeek et al., 2001; Schade et al., 1993). Another appetite stimulating hormone orexin promotes wakefulness while stimulated by ghrelin and enhances sleep when inhibited by leptin (Sakurai, 2007) and orexigenic neuron activation is believed to be under circadian control (Marston et al., 2008). Short neuropeptide F (sNPF) signaling plays a role in feeding and sleep (Lee et al., 2004; Shang et al., 2013). sNPF receptor (sNPFR) signaling in Insulin Producing Cells (IPCs) regulate the expression of Drosophila Insulin like peptides (DILPS) and metabolism in Drosophila (Lee et al., 2008; Fadda et al., 2019). sNPF is expressed in circadian clock small ventral lateral nurons (s-LNvs) and dorsal lateral neurons (LNds) regulates sleep in *Drosophila* (Shang et al., 2013).

Feeding and sleep wake cycles are interconnected behaviors and the optimum balance between them is disrupted under nutrient deprivation. Starvation suppresses sleep by promoting hyperactivity in *Drosophila* and this behavioral change in response to nutrient unavailability is governed by diverse neuropeptides such as allatostatin, octopamine, adipokinetic hormone, glucagon and insulin (Chen et al., 2016; G. Lee and Park, 2004; Yang et al., 2015; Yu et al., 2016). Sustained increase in locomotor activity under starvation condition partly resembles foraging behavior (Yang et al., 2015). Feeding behavior appears to be under multiple controls including the gustatory, olfactory and circadian neural circuits (Xu et al., 2008; Lin et al., 2019). Significant progress has been made in understanding the circadian and neuronal basis of feeding rhythm and starvation-mediated sleep changes in *Drosophila* (Xu et al., 2008; Keene et al., 2010). However it remains unclear whether the circadian clock conveys the time of the day information to coherently coordinate this starvation-mediated behavioral change with metabolism through interlinked neuronal pathways or via independent neural circuitry.

In the present study, we aimed to elucidate the role of Short neuropeptide F in triglyceride metabolism, starvation-mediated enhanced activity and starvation resistance in *Drosophila*. The results showed that the circadian clock regulates the transcription of *snpf* gene. sNPF expressed in circadian clock neurons delays the starvation-mediated hyperactivity response, whereas sNPF receptor expressed in IPCs advances the starvation-mediated hyperactivity response in *Drosophila*. sNPF expressed in clock neurons enhance the triglyceride level and starvation resistance whereas SNP receptors expressed in the IPCs reduce the starvation resistance suggesting the differential role of neuropeptide sNPF in clock neurons and IPCs to regulate the starvation mediated locomotor activity in accordance with the nutrient availability in *Drosophila*.

## Materials and Methods

### Fly stocks and maintenance

Following fly lines were used in this experiment: *w^1118^* (BDSC #5905), *tim-Gal4 and per^o^* (SheebaVasu Laboratory), *dilp2-Gal4, UAS-snpf^RNAi^* and UAS-*snpfR^RNAi^* (Kweon Yu Laboratory). All fly stocks were maintained in the *Drosophila* growth chamber (MIR-154, Panasonic, Japan) at 25° C temperature, 75 ± 5% humidity and 12:12 hour (h) light-dark (LD) cycle where lights came on at Zeitgeber Time 00 (ZT 00) and went off at ZT12.

### Measurement of mRNA levels

To assess the mRNA transcription profiles of *snpf*, three replicates were used for each time point. Each replicate contained 30 heads from 2-3 day old *w^1118^* male flies. Total RNA was extracted using QIAGEN RNAeasy Plus Mini kit; DNAase digestion was performed using QIAGEN RNase–free DNase. cDNA was synthesized using SuperScript™ III First-Strand Synthesis System (Invitrogen, Cat. No. 18080051) and real-time PCR was performed using Bio-Rad CFX96TM with the cDNA template, Power SYBR^®^ Green PCR Master Mix (Cat. # 4368702) from ThermoFisher Scientific to check temporal transcription profile of *snpf*. mRNA levels were measured under LD cycle and constant darkness (DD). The molecular oscillation of mRNA levels was determined by normalizing the mRNA level of the gene of interest with the mRNA level of *rp49* at each time point. Knockdown efficiency of *tim-Gal4* > *snpf^RNAi^* was validated by qPCR. We used the following primers for qPCR. *snpf* forward primer 5’-CCCGAAAACTTTTAGACTCA-3’ and reverse primer 5’-TTTTCAAACATTTCCATCG-3’, *rp49* forward primer 5’-GCTAAGCTGTCGCACAAA-3’ and reverse primer 5’-TCCGGTGGGCAGCATGTG-3’.

### Locomotor Activity assay

For activity/rest assay, freshly emerged male flies were pre-entrained for two days in the LD cycle at 25 °C and a humidity of 75 ± 5% inside the incubator (MIR-154, Panasonic, Japan). Two-day-old virgin males were individually introduced into activity tubes. Flies were provided with standard cornmeal medium and their activity/rest behavior was recorded using Drosophila Activity Monitors (DAM, Trikinetics, USA). The infrared beam that passes through the middle of the activity tubes count the movement of the flies. Following acclimatization in activity tubes for 12 hours, flies were transferred to 1% agar medium or to standard cornmeal medium in glass tubes at ZT12 (during the onset of dark phase) to provide starved or fed conditions, respectively, and locomotor activity was recorded for 36 hours at 25 °C. Each experiment was performed with 27-32 flies for each genotype and DAM readings were collected in every 1-minute interval and the activity counts/ hour is plotted in the locomotor activity waveform. Flies died within 24 hour of starvation were excluded and flies survived for 24-36 hours of starvation were used for the analysis. Change in activity under starvation was calculated as [(Activity under starvation-Activity under fed) /Activity under fed condition] x 100. Separate sets of flies were used for recording the locomotor activity under fed and starved flies condition. Hence the change in activity under starvation condition was calculated as [(Activity of individual fly under starvation-Average activity of all the flies under fed) / Average activity of all the flies under fed] x 100.

### Triglyceride assay

This assay comprised of three replicates and each replicate containing five male flies under fed condition was homogenized in homogenization buffer (0.05% Tween 20). The homogenate was incubated at 70°C for 5 minutes to inactivate the enzymes and centrifuged at 14000 rpm for 3 minutes and the supernatant was collected into a sample tube. Sigma serum triglyceride determination kit TR0100 was used to estimate the triglyceride level in the sample by measuring absorbance at 540 nm using TECAN Infinite M200 pro-multimode plate reader. The triglyceride content was normalized to the total protein content of the flies estimated by Quick Start™ Bradford 1X Dye Reagent (BioRad, Cat. # 500-0205). Freshly emerged flies feed less and possess larval triglyceride energy storage. Larval fat cells persist only up to about two days and are replaced by adult fat cells. Hence assays were carried out at ZT 12 in fed two-day-old male flies to estimate the triglyceride levels in adult flies.

### Starvation resistance Assay

Starvation resistance assay comprised of six replicates with each replicate vial containing 10-15 flies for each genotype. Five-day-old male flies were kept in the 1% agar medium vials. The number of dead flies was recorded at every two-hour interval until all the flies died.

### Statistical analysis

Cosinor analysis was implemented in MATLAB-R2016a to test for rhythmicity, to estimate the peak phase and the amplitude (peak/trough ratio) of relative mRNA expression of *snpf* (Nelson et al., 1979). The waveform of activity/rest rhythm, day time activity, nighttime activity and change in locomotor activity were analyzed by using ANOVA followed by Tukey’s post hoc multiple comparisons. Log-rank test was used to compare and analyze the survivorship curves under starvation. TGA levels was analyzed by Student’s t-test. The statistical analyses were implemented on Statistica for Windows Release 5.0 B (Statsoft, 1995). Error bars in all the graphs represent standard error of the mean (SEM).

## Results

### *snpf* transcript exhibit circadian oscillation

To assess whether the circadian clock regulates *snpf* transcript oscillation, the transcription profile of *snpf* was estimated in *w^1118^* flies under LD and under constant darkness (DD). The *snpf* mRNA level increased during light phase and decreased during the dark phase resulting in a significant daily oscillation with a peak at ZT 8.1h under LD (Cosinor analysis, *p*<0.001) (Fig. 1A). The *snpf* transcriptional oscillation persisted under DD with a peak at CT 8.5h (Cosinor analysis, *p*<0.002). However, the amplitude (peak/trough ratio) of the oscillation decreased under DD compared to that obtained under LD (Fig. 1B). To confirm the role of circadian clock in *snpf* transcript oscillation, further studies were carried out in *per^o^* flies under LD. *snpf* transcript level in *per^o^* flies did not exhibit any significant rhythmicity under LD (cosinor analysis, *p*> 0.05) (Fig. 1C). These results suggest that the transcriptional oscillation of *snpf* mRNA is endogenously controlled by the clock.

**Figure 1.**
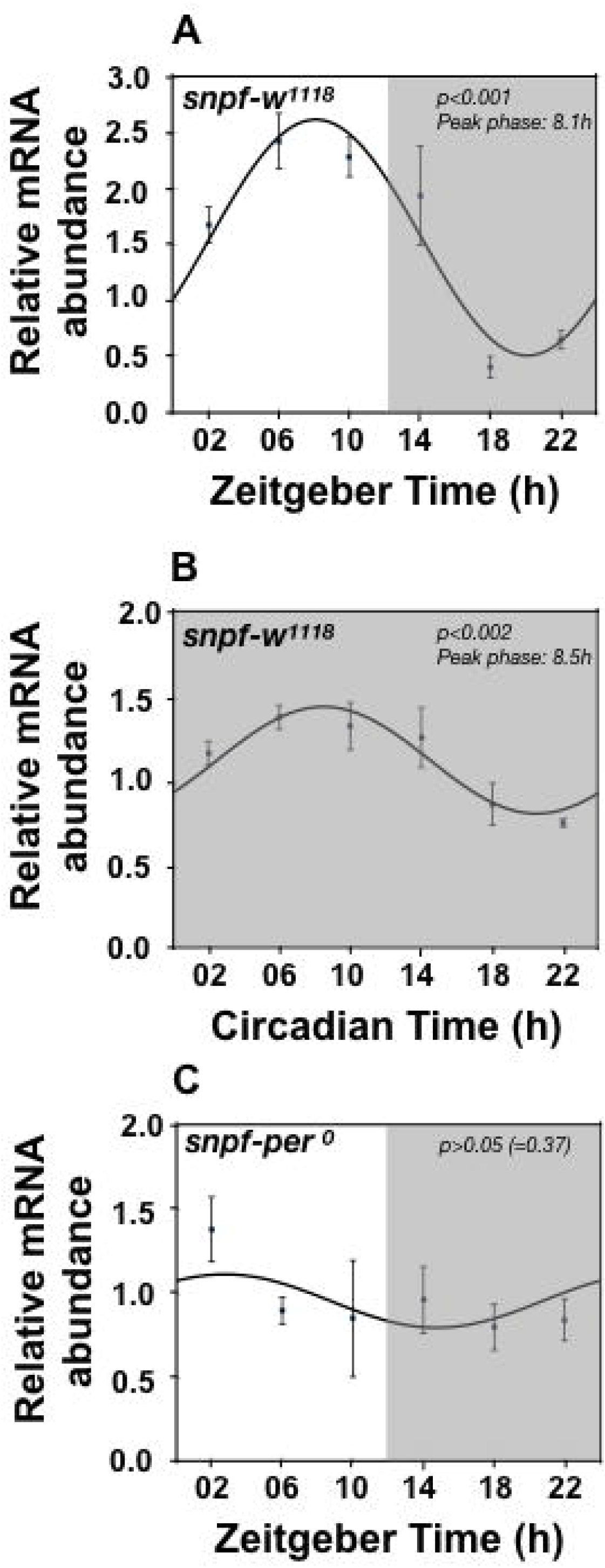
Transcription of sNPF mRNA is clock controlled. (A) Transcription profile of *snpf* mRNA in *w^1118^* flies under 12:12 h light-dark cycle (LD). White and grey shaded bars represent light and dark phase, respectively, under LD. (Cosinor analysis, *p*<0.001) (B) Transcription profile of *snpf* mRNA in *w^1118^* flies constant darkness (DD). *snpf* transcript exhibited a significant daily oscillation with a peak at ZT 8.1h under LD and the oscillation persisted under DD with a peak at CT 8.5h. (Cosinor analysis, *p*<0.002) (C) Transcription profile of *snpf* mRNA in *per^0^* flies under 12:12 h light-dark cycle (LD). Three replicates were used for each time point. Each replicate contained 30 heads from 2-3 day old *w^1118^* male flies. Error bars represent mean ± SEM.

### sNPF expressed in circadian clock neurons suppress starvation-mediated hyperactivity

To assess the role of sNPF expressing circadian clock neurons in starvation-mediated hyperactivity, we expressed *UAS-snpf^RNAi^* under the *tim-Gal4* driver (*tim-Gal4* > *snpf^RNAi^*) and recorded the locomotor activity under fed and starved LD conditions for 36 h. *tim-Gal4* > *w^1118^*, *w^1118^* > *UAS-snpf^RNAi^* control flies and the *tim-Gal4* > *snpf^RNAi^* flies exhibited increased activity under starved condition compared to what was observed under fed condition (*tim-Gal4* > *w^1118^* flies; ANOVA, *p*<0.0001; *w^1118^* > *UAS-snpf^RNAi^*; ANOVA, *p*<0.0001; *tim-Gal4* > *snpf^RNAi^* flies; ANOVA, *p*<0.0001) (Fig. 2A-C). ANOVA followed by post-hoc multiple comparisons using Tukey’s test on activity data showed that starved *tim-Gal4* > *w^1118^* flies enhanced the activity at 25-27h of starvation (Fig. 2A) resulting in a significant increase in the total activity second dark phase (25-36h) (ANOVA, *p*<0.0001) and during the light phase (13-24h) (ANOVA, *p*<0.0001) compared to those observed under fed condition (Fig. E-F). *w^1118^* > *snpf^RNAi^* flies enhanced the activity at 1-2h, 16-19h, 25-27h of and 32h starvation. This resulted in a significant increase in the total activity of *w^1118^* > *snpf^RNAi^* flies during the first dark phase (1-12h) (ANOVA, *p*<0.0001), light phase (13-24h) (ANOVA, *p*<0.0001) and second dark phase (25-36h) (ANOVA, *p*<0.0001) under starvation compared to those observed under fed condition (Fig. 2B, D-F). *tim-Gal4* > *snpf^RNAi^* flies exhibited increased activity at 15h, 25-31h and 33h of starvation resulting in a significant increase in the total activity during the light phase (13-24h) (ANOVA, *p*<0.0001) and second dark phase (25-36h) (ANOVA, *p*<0.0001) under starvation compared to those observed under fed condition (Fig. 2C, E-F). Comparison of change in activity between controls and *tim-Gal4* > *snpf^RNAi^* flies under starved vs fed condition revealed that *tim-Gal4* > *snpf^RNAi^* flies exhibited significant increase in activity after 24 hours of starvation (25-36h) than *tim-Gal4* > *w^1118^* (ANOVA, p<0.0001) and *w^1118^* > *snpf^RNAi^* (ANOVA, p<0.0001) controls (Fig. 2G). These results suggest that sNPF in circadian clock neurons delays the starvation-mediated increased locomotor activity response after 24h of starvation in *Drosophila*.

**Figure 2.**
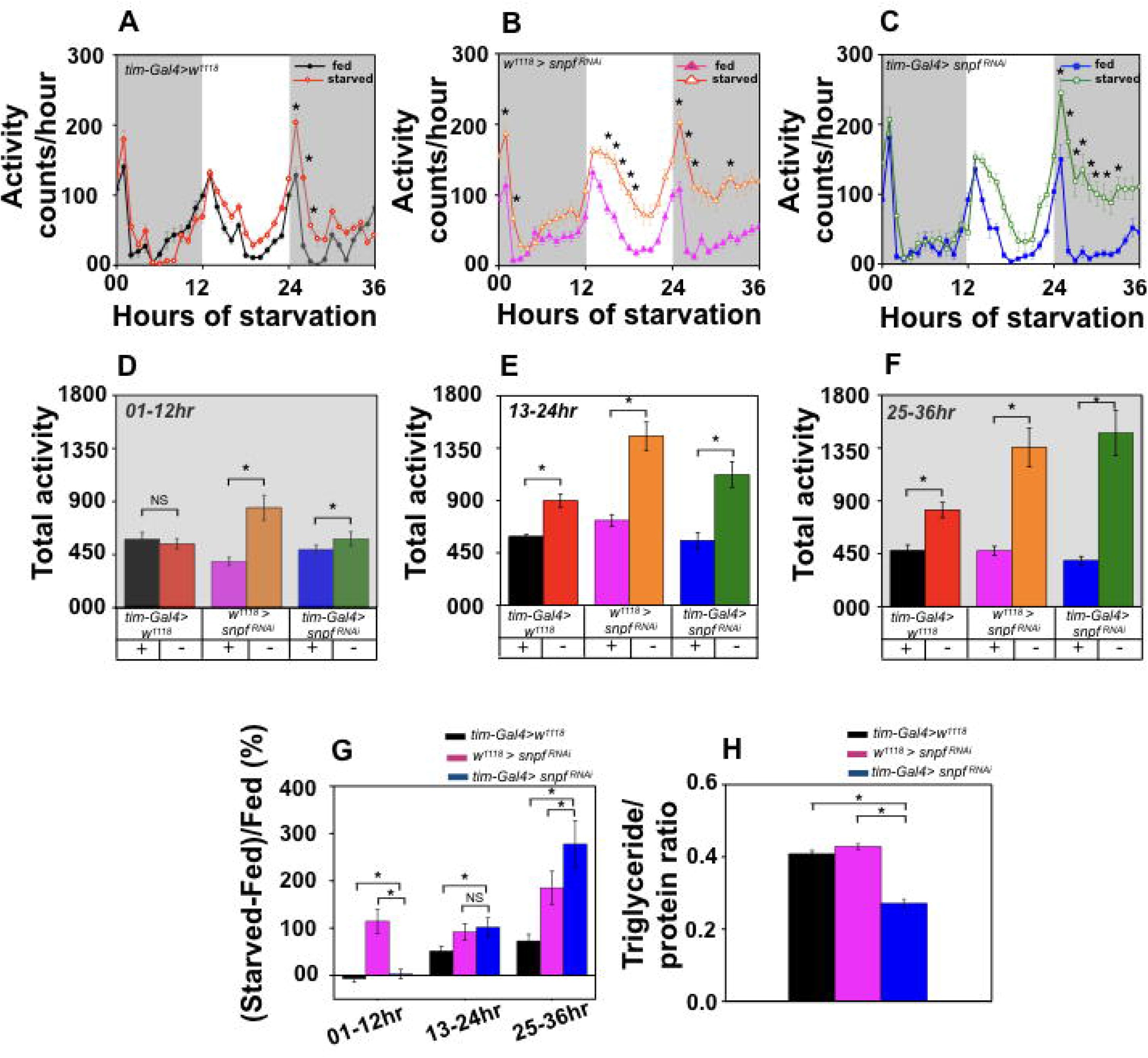
Role of sNPF expressing circadian clock neurons in starvation-mediated hyperactivity. (A-C) Average locomotor activity counts/hour of *tim-Gal4* > *w^1118^*, *w^1118^* > *snpf^RNAi^* and *tim-Gal4* > *UAS-snpf^RNAi^* (*tim-Gal4* > *snpf^RNAi^*) flies assayed with ad libitum food and under starvation for 36 h. Flies were transferred to fresh food or to 1% agar medium at the onset of dark phase to provide the fed and starved condition respectively. Activity counts per hour is plotted along the *y*-axis and hours of starvation on *x*-axis. Gray shaded bars indicate dark phase under LD for 36 h time window. Control flies *tim-Gal4* > *w^1118^* (*n*= 27) (ANOVA, *p*<0.0001), *w^**1118**^* > *snpf^RNAi^* (*n*= 30) (ANOVA, *p*<0.0001) and *tim-Gal4* > *snpf^RNAi^* (*n*= 27) (ANOVA, *p*<0.0001) exhibited enhanced locomotor activity under starvation condition compared to fed controls. (D-F) Total activity of *tim-Gal4* > *w^1118^*, *w^**1118**^* > *snpf^RNAi^* and *tim-Gal4* > *snpf^RNAi^* flies during the first dark phase (1-12h), light phase (13-24h) and second dark phase (25-36h) under fed and starved conditions. (G) Change in activity under starved condition compared to the fed condition. Starved experimental flies (*tim-Gal4* > *snpf^RNAi^*) exhibited a significant increase in the activity after 24 h of starvation compared to both *tim-Gal4* > *w^1118^* (ANOVA, *p*<0.0001) and *w^1118^* > *snpf^RNAi^* (ANOVA, *p*<0.0001) controls. (H) Flies expressing *snpf^RNAi^* in clock neurons exhibit reduced triglyceride levels on day 2 compared to both *tim-Gal4* > *w^1118^* (t-test, *p*<0.0001) and *w^1118^* >*snpf^RNAi^* controls (t-test, *p*<0.0001) (*N*=3; *n*= 5). The asterisks on locomotor activity waveform and the vertical bar graphs indicate that the values are significantly different. Error bars represent mean ± SEM. For further explanations see text.

We further assessed the role of sNPF in regulating the triglyceride levels and starvation resistance in flies expressing *UAS-snpf^RNAi^* under the *tim-Gal4* driver. *tim-Gal4* > *snpf^RNAi^* flies exhibited a significant decrease in the triglyceride level compared to both *tim-Gal4* > *w^**1118**^* (t-test, *p*<0.0001) and *w^**1118**^* > *snpf^RNAi^* controls (t-test, *p*<0.0001) (Fig. 2H). In agreement with the enhanced triglyceride level, *tim-Gal4* > *snpf^RNAi^* flies exhibited a significant decrease in the starvation resistance compared to both *tim-Gal4* > *w^1118^* (log-rank test, *p*<0.0001) and *w^1118^* > *snpf^RNAi^* controls (log-rank test, *p*<0.0001) (Fig. S1A). These results suggest that knock down of *snpf* in circadian clock neurons reduces the triglyceride level and starvation resistance in *Drosophila*.

### sNPFR expressed in IPC promotes starvation-mediated hyperactivity

To test the role of sNPFR expressed in IPCs on starvation-mediated hyperactivity, we expressed the *UAS-snpfR^RNAi^* under the *dilp2-Gal4* driver (*dilp2-Gal4* > *snpfR^RNAi^*) and recorded the locomotor activity of these flies under fed and starved condition for 36 h. *dilp2-Gal4* > *w^1118^*, *w^1118^* > *snpfR^RNAi^* control flies and the *dilp2-Gal4* > *snpfR^RNAi^* flies exhibited increased activity under starved condition compared to what was observed under fed condition (*dilp2-Gal4* > *w^1118^* flies; ANOVA, *p*<0.0001; *w^1118^* > *snpfR^RNAi^*; ANOVA, *p*<0.0001; *dilp2-Gal4* > *snpfR^RNAi^* flies; ANOVA, *p*<0.0001) (Fig. 3A-C). ANOVA followed by post-hoc multiple comparisons using Tukey’s test on activity data showed that starved *dlip2-Gal4* > *w^1118^* flies enhanced the activity at 1h, 13-17h, 23-27h and 30-32h of starvation resulting in a significant increase in the total activity during the first dark phase (1-12h) (ANOVA, *p*<0.005), light phase (13-24h) (ANOVA, *p*<0.0001) and second dark phase (25-36h) (ANOVA, *p*<0.0001) compared to those observed under fed condition (Fig. 3A, D-F). *w^**1118**^* > *snpfR^RNAi^* flies enhanced the activity at 2h, 16-17h, 25-27h and 29-31h of starvation. This resulted in a significant increase in the total activity of *w^1118^* > *snpfR^RNAi^* flies during the light phase (13-24h) (ANOVA, *p*<0.0001) and second dark phase (25-36h) (ANOVA, *p*<0.0001) compared to those observed under fed condition (Fig. 3B, E-F). *dilp2-Gal4* > *snpfR^RNAi^* flies exhibited increased activity at 14-16h and 23-28h of starvation resulting in a significant increase in the total activity only during second dark phase (25-36h) (ANOVA, *p*<0.001) and under starvation compared to those observed under fed condition (Fig. 3C, F). Starvation did not enhance the activity in experimental flies (*dilp2-Gal4* > *snpfR^RNAi^*) during the first 24 hours of starvation compared to fed flies (Fig. 3C-E). Comparison of change in activity between controls and *dilp2-Gal4* > *snpfR^RNAi^* flies under starved vs fed condition revealed that *dilp2-Gal4* > *snpfR^RNAi^* flies exhibited decrease in activity than controls after 24 hours of starvation (25-36h) than *dilp2-Gal4* > *w^1118^*(ANOVA, *p*<0.0001) and *w^1118^* > *snpfR^RNAi^* (ANOVA, *p*<0.03) control flies (Fig. 3G). These results suggest knock down of the *snpfR* expressed in IPCs delays the starvation-mediated hyperactivity in Drosophila.

**Figure 3.**
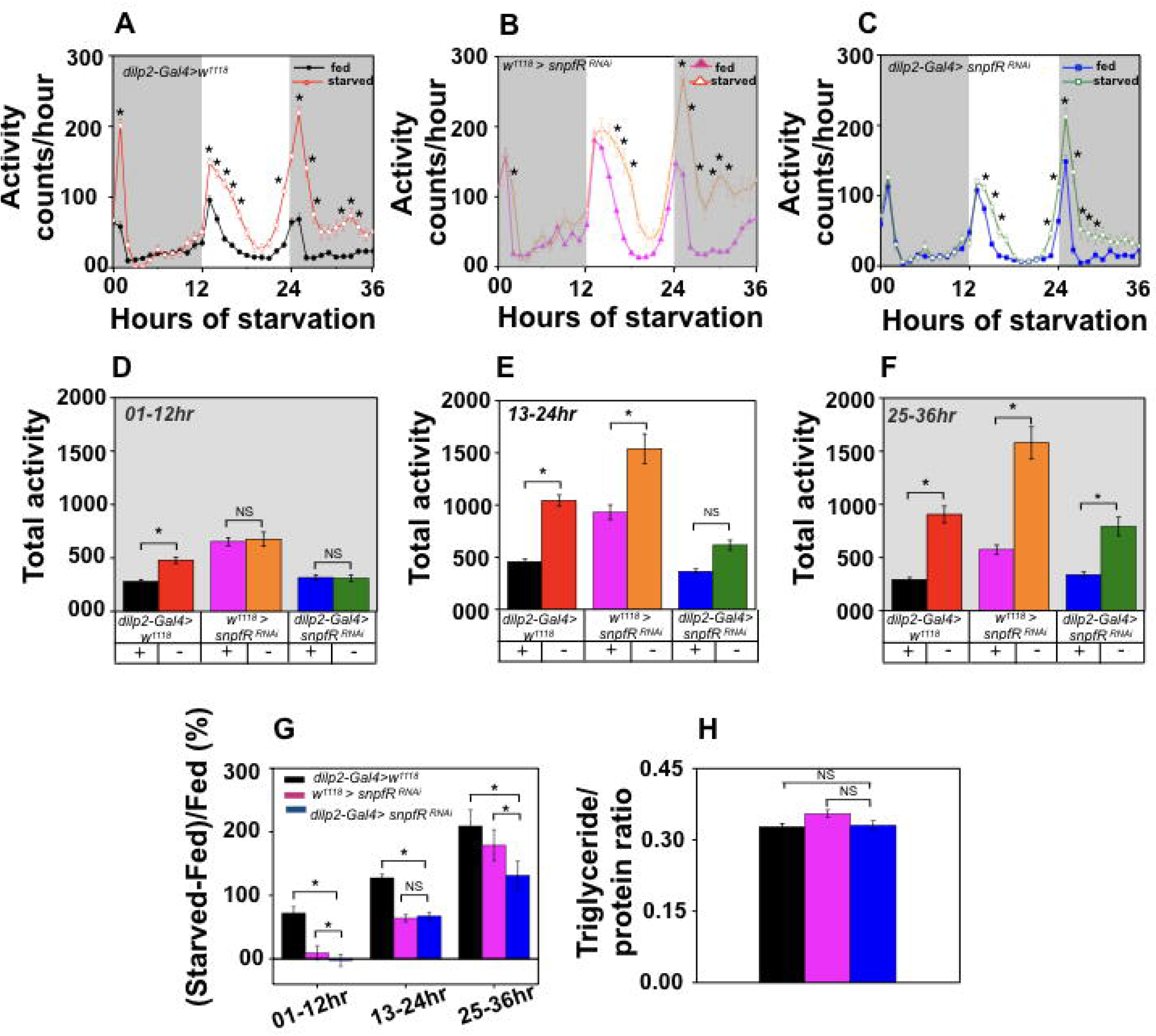
Role of SNPF receptors expressed in IPC in regulating starvation-mediated hyperactivity. (A-C) Average activity counts/hour of *dilp2-Gal4* > *w^1118^*, *w^1118^* > *snpfR^RNAi^* and *dilp2-Gal4* > *UAS-snpfR^RNAi^* (*dilp2-Gal4* > *snpfR^RNAi^*) flies assayed under the fed and starved condition.. Control flies *dilp2-Gal4* > *w^1118^* (*n*= 29) (ANOVA, *p*<0.0001), *w^**1118**^* > *snpfR^RNAi^* (*n*= 31) (ANOVA, *p*<0.0001) and *dilp2-Gal4* > *snpfR^RNAi^* (*n*= 30) (ANOVA, *p*<0.0001) exhibited enhanced locomotor activity under starvation condition compared to fed controls. (D-F) Total activity of *dilp2-Gal4* > *w^1118^*, *w^111 8^* > *snpfR^RNAi^* and *dilp2-Gal4* > *snpfR^RNAi^* flies during first dark phase (1-12h), light phase (13-24h) and second dark phase (25-36h) under fed and starved conditions. (G) Change in activity under starved condition compared to the fed condition. Experimental flies (*dilp2-Gal4* > *snpfR^RNAi^* exhibited reduction in the activity after 24 hours of starvation compared to both *dilp2-Gal4* > *w^1118^* (ANOVA, *p*<0.0001) and *w^1118^* > *snpfR^RNAi^* (ANOVA, *p*<0.03) controls. (H). Triglyceride levels of two day old *dilp2-Gal4* > *w^1118^*, *w^1118^* > *snpfR^RNAi^* and *dilp2-Gal4* > *snpfR^RNAi^* flies (*N*=3; *n*= 5). Other details are the same as in Fig 2.

We also assessed whether *snpf*R expressed in the IPCs regulate the triglyceride levels and starvation resistance in flies by expressing *UAS-snpfR^RNAi^* under the *dilp2-Gal4* driver. *dilp2-Gal4* > *snpfR^RNAi^* flies did not exhibit any significant difference in the triglyceride level compared to both *dilp2-Gal4* > *w^**1118**^* (t-test, *p*>0.05) and *w^1118^* > *snpfR^RNAi^* controls (t-test, *p*>0.05) (Fig. 3H). However, *dilp2-Gal4* > *snpfR^RNAi^* flies exhibited a significant increase in the starvation resistance compared to both *dilp2-Gal4* > *w^1118^* (log-rank test, *p*<0.0001) and *w^1118^* > *snpfR^RNAi^* controls (log-rank test, *p*<0.0001) (Fig. S1B). These results suggest that knock down of *snpfR* in IPCs increases the starvation resistance in *Drosophila*.

## Discussion

Behavior and metabolism are functionally intertwined processes that modulate each other. However, it remains elusive how the circadian clock governs the major clock mediated behavioral output locomotor activity rhythm, in synchrony with the metabolic status of an organism. In the present study, we aimed to examine the specific role of circadian neuropeptide, i.e., sNPF in lipid metabolism and starvation-mediated behavioral changes in *Drosophila*. The results of the present study showed that *snpf* transcript exhibited a circadian oscillation in the fly head. sNPF protein is expressed in a diverse subset of neurons in the adult fly brain including olfactory receptor neurons, various small interneurons, kenyon cells of the mushroom bodies and in the circadian clock neurons (Nässel., 2008). Among the ~150 clock neurons clustered in the fly brain, two dorsal lateral neurons (LNds) and PDF expressing small ventral lateral neurons (s-LNvs) express sNPF (Johard et al., 2009). Based on the previous findings, mRNA cycling is essential for the circadian clock neuron function including the neuronal firing or electrical excitability (Kula-Eversole et al., 2010; Shang et al., 2013). Hence it is quite likely that the *snpf* transcript oscillation is more apparent in circadian clock neurons than other sNPF expressing diverse neuronal subsets. Our results showed that *snpf* transcript level did not exhibit any significant rhythmicity *period* null mutants confirming the endogenous regulation of *snpf* transcript oscillation. Most of the previously identified cycling head transcripts are tissue or cell-specific in the mammalian system (Storch et al., 2002). Ubiquitous mRNA cycling is relatively rare. Further studies are required to confirm whether sNPF oscillation is indeed restricted to clock neurons or if it also oscillates in non-clock neurons.

In agreement with the previous studies, we showed that starvation enhances locomotor activity in *Drosophila* (Lee and Park 2015; Yang et al., 2015). Given that starvation-induced hyperactivity and food intake are mutually inhibitory (Chen et al., 2015; Yang et al., 2015), it is of great interest to understand how these two behaviors are dynamically regulated by the central and peripheral neural circuits. sNPF expression in a large population of diverse neurons indicates the multiple discrete functions of this neuropeptide (Nässel., 2008). So far studies demonstrated that sNPF regulates feeding and its expression in the fan shaped body might be involved in fine-tuning the locomotor activity level (Lee et al., 2004; Kahsai et al., 2010). Expression of sNPF and sNPF receptor in the mushroom body and clock neurons regulate sleep in *Drosophila* (Chen et al., 2013; Shang et al., 2013). sNPF released from s-LNvs and LNds governs the Dorsal neuron 1 (DN1) late night Ca^2+^ activation and the Ca^2+^ dynamics in DN1 neurons are intricately linked to behavioural rhythm (Liang et al., 2017). Downregulation of sNPF in all pacemaker neurons increased nighttime locomotor activity under fed condition (Liang et al., 2017). The results of our present studies showed that knockdown of *snpf* in circadian clock neurons advances the starvation-mediated hyperactivity during the second dark phase, suggesting the role of sNPF expressing clock neurons in delaying hyperactivity under starvation. In the present study starvation condition was provided at the onset of dark phase (ZT 12) and flies expressing *snpf* RNAi in clock neurons exhibited enhanced starvation mediated hyperactivity after 24 hours of starvation during the second dark phase. sNPF is a sleep promoting molecule (Shang et al., 2013) and it regulates the Ca^2+^ activation during the nighttime in DN1 neurons (Liang et al., 2017) that possibly modulates the starvation-induced hyperactivity during the darkness. In the present study starvation condition was provided at the onset of dark phase (ZT 12), further studies are required on *tim-Gal4* > *snpf^RNAi^* flies with the starvation starting at the onset of light phase (ZT00) to elucidate whether increased starvation-mediated hyperactivity observed after 24hr of starvation is restricted to the second dark phase or it would persist even under light phase after 24hr of starvation. A population of dorsally located *clk*-expressing neurons has been implicated in promoting sleep during starvation (Keene et al., 2010). Among the clock neurons, PDF positive morning cell s-LNvs and PDF negative evening cells LNds express sNPF (Johard et al., 2009). Morning cell specific downregulation of sNPF impaired morning anticipatory activity under fed condition. sNPF from both M cells and E cells are required for the Ca2+ rhythmicity, its phase and behavioral output (Liang et al., 2017). Further studies are required to confirm the specific role of sNPF expressing PDF positive s-LNvs and PDF negative LNds in starvation-mediated hyperactivity.

Although sNPF predominantly acts as an inhibitory modulator in neuronal networks (Vecsey et al., 2014) sNPFR expressed in olfactory receptor neurons has excitatory effect and promotes starvation induced food search behavior (Root et al., 2011). Knockdown of *snpfR* in IPCs in an adult stage-specific manner does not affect the starvation-mediated sleep loss in *Drosophila* (Shang et al., 2013), whereas the results of our present study showed that constitutive knockdown of *snpfR* in IPCs suppress starvation-mediated hyperactivity after 24 hours of starvation. While sNPFR expressed in IPCs promote starvation-mediated hyperactivity, sNPF expressed in circadian clock neurons had an inhibitory effect on this starvation mediated behavioral phenotype.

Knock down of sNPF in sNPF producing dorso lateral peptidergic (DLP) neurons increased the starvation resistance and triglyceride levels in *Drosophila* (Kapan et al., 2012). In contrast, we observed that *tim-Gal4* > *snpf^RNAi^* flies exhibit decrease in the triglyceride level and starvation resistance whereas knock down of *snpfR* expressed in IPCs enhanced only the starvation resistance in *Drosophila* indicating the differential function of *snpf* expressing clock neurons and *snpR* expressing IPCs in regulating the starvation resistance.

Taken together, our present study showed that transcriptional oscillation of *snpf* mRNA is endogenously controlled by the circadian clock and sNPF expressed in circadian clock neuron delays the starvation-mediated hyperactivity response. sNPF might promote starvation induced locomotor activity partially through sNPF receptors expressed in the IPCs. The sNPF appears to be a central neuropeptidergic pathway in regulating starvation mediated hyperactivity, triglyceride metabolism and starvation resistance in *Drosophila*.

## Acknowledgements

We thank Sabira A for the experimental assistance. We thank Jishy Varghese for the valuable suggestions and for providing the infrastructure facility to carry out some of the experiments, Kenji Tomioka for the valuable suggestions, SheebaVasu for providing the fly lines and Nikhil K L for helping with the statistical analysis.

## Author Contributions

Conceived and designed the experiments: NNK AG AEK HP. Performed the experiments: AG AEK HP SRS. Analyzed the data: AG AEK HP. Contributed reagents/materials: NNK. Wrote the paper: AG NNK.

## Supporting information

**Figure S1.** (A) Percentage survivorship of 5 day old flies expressing *snpf^RNAi^* in clock neurons under starvation condition. Five-day-old flies expressing *snpf^RNAi^* in clock neurons exhibit reduced starvation resistance compared to the control flies (log-rank test, *p*<0.0001). (B) Percentage survivorship of 5 day old flies expressing *snpfR^RNAi^* in IPCs under starvation condition. Flies expressing *snpfR^RNAi^* in IPCs exhibit reduced starvation resistance compared to the control flies (log-rank test, *p*<0.0001).

