## Supplementary figures and images for "Short Neuropeptide F regulates the starvation mediated enhanced locomotor Activity in *Drosophila*"

### Supplementary Figure S1

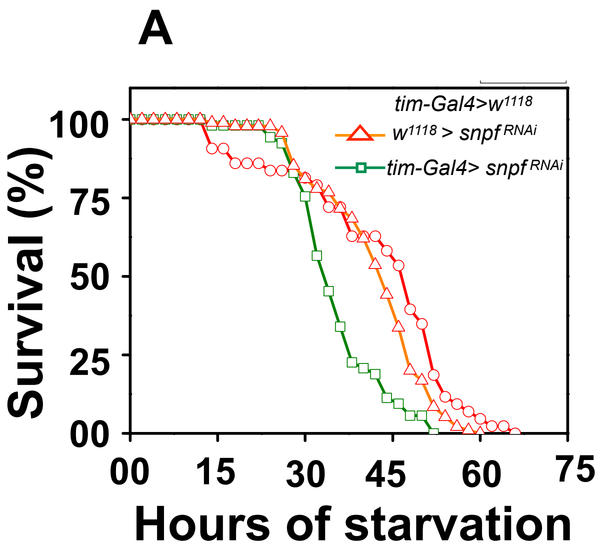

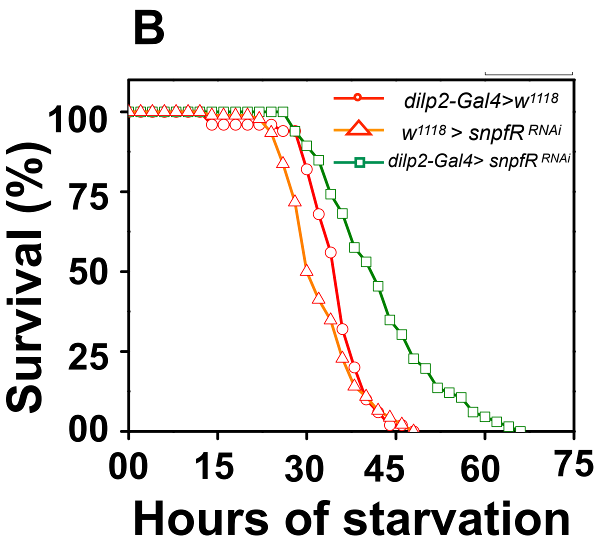
